# PP2B-dependent cerebellar plasticity sets the amplitude of an innate reflex during juvenile development

**DOI:** 10.1101/2024.01.24.576981

**Authors:** Bin Wu, Laura Post, Zhanmin Lin, Martijn Schonewille

## Abstract

Throughout life, the cerebellum plays a central role in the coordination and optimization of movements, using cellular plasticity to adapt a range of behaviors. If these plasticity processes establish a fixed setpoint during development, or continuously adjust behaviors throughout life, is currently unclear. Here, by spatiotemporally manipulating the activity of protein phosphatase 2B (PP2B), an enzyme critical for cerebellar plasticity, we examined the consequences of disrupted plasticity on the performance and adaptation of the vestibulo-ocular reflex (VOR). We find that, in contrast to Purkinje cell specific deletion starting early postnatally, acute pharmacological as well as adult-onset genetic deletion of PP2B affects all forms of VOR adaptation, but not the level of VOR itself. Next, we show that Purkinje cell-specific genetic deletion of PP2B in juvenile mice leads to a progressive loss of the protein PP2B and a concurrent change in the VOR, in addition to the loss of adaptive abilities. Finally, re-expressing PP2B in adult mice that lack PP2B expression from early in development, rescues VOR adaptation, but does not affect the performance of the reflex. Together, our results indicate that chronic or acute, genetic or pharmacological block of PP2B disrupts the adaptation of the VOR. In contrast, only the absence of plasticity during cerebellar development affects the setpoint of VOR, an effect that cannot be corrected after maturation of the cerebellum. These findings suggest that cerebellar plasticity influences behavior in two ways, through direct control of behavioral adaptation and via long-term effects initiated in the juvenile period.

**Significance Statement:** Early damage to motor adaptation structures, such as the cerebellum, has been linked to neurodevelopmental disorders persisting into adulthood. Understanding these long-term effects requires disentangling the persistent, long-term effects of disrupted development from the acute, ongoing effects directly caused by the continuous presence of the disruption. Here, we demonstrate that disruptions during early development affect both basal level and adaptation, whereas late, adult-onset disruption of cerebellar plasticity only affects the ability to adapt, not the setpoint of an innate reflex. Our findings that specifically the absence of plasticity during cerebellar development affects the setpoint of VOR, which cannot be corrected by re-instating plasticity after maturation of the cerebellum, supports the concept of a sensitive developmental period for setting innate reflexes.

## Introduction

The cerebellum plays a crucial role in motor control and adaptation by integrating sensory and motor information (De Zeeuw CI and Yeo CH, 2005; Ito M, 2002). Formation of the adult cerebellar cyto-architecture, through neurogenesis, migration, morphological maturation and circuitry finetuning, is initiated prenatally, but completed in the postnatal period, in both man and mice (Beekhof GC et al., 2021; Marzban H et al., 2014). In line with the fact that it is well-conserved throughout phylogenesis and that it develops relatively late during ontogeny, the cerebellum has a highly organized anatomical structure that can subserve the control of various behavioral and cognitive functions (Bayer SA et al., 1993; Hatten ME and Heintz N, 1995; Larsell O, 1952; Sillitoe RV et al., 2005). Accordingly, the cerebellum is often used as a model system for investigating the interactions between development and motor performance and motor learning (Manto MU and Jissendi P, 2012; Martinez S, 2014). Peri- and postnatal damage of the cerebellum has been linked to neurodevelopmental disorders with functional deficits that can persist well into adulthood, highlighting the possibility that the cerebellum may contribute to more wide-spread neurotypical brain maturation (Leto K et al., 2016; Wang R et al., 2018; Wang SS et al., 2014). Understanding these long-term effects requires disentangling the persistent, long-term effects of disrupted development from the acute, ongoing effects directly caused by the disruption. Based on the premise that cerebellar damage early in life can be linked to problems at later stages, we hypothesize that there is a differential effect of early versus late disruption of cerebellar function, with more basal functions affected by developmental disruption.

To quantitatively evaluate the influence of postnatal development on both motor performance and motor learning across the entire life span, it will be advantageous to study a cerebellum-dependent behavior that allows for measuring both behavioral components, that shows minimal variation among subjects, and that can be reliably recorded over a wide range of ages. Compensatory eye movements perfectly meet these criteria as they consist of adaptable reflexes induced by vestibular and optokinetic inputs (Faulstich BM et al., 2004; Katoh A et al., 1998; Miles FA and Lisberger SG, 1981; Stahl JS, 2004). Compensatory eye movement reflexes and their adaptation can be rapidly probed in a highly reproducible manner from birth in humans (Weissman BM et al., 1989) and shortly after eye opening in mice (Beekhof GC, et al., 2021; Faulstich BM, et al., 2004), making it an ideal system to study the acute and long-term effects of interventions at various stages in life. The expression of protein phosphatase 2B, also known as calcineurin, starts early postnatally (Lin Z et al., 2021) and is critical for cerebellar plasticity, including parallel fiber to Purkinje cell long-term potentiation (PF-PC LTP) and plasticity of intrinsic excitability (IP) of Purkinje cells (Belmeguenai A and Hansel C, 2005; Belmeguenai A et al., 2010; Schonewille M et al., 2010). Mice harboring a Purkinje cell specific knockout of PP2B (L7-PP2B KO) from early on display severe deficits in a broad range of cerebellum-dependent behaviors including performance and adaptation of compensatory eye movements (Lefort JM et al., 2019; Rahmati N et al., 2014; Schonewille M, et al., 2010; Vinueza Veloz MF et al., 2015), but whether these deficits are due to the absence of PP2B at the adult stage or to disrupted cerebellar development, remains unclear.

Here, using comparative behavioral analysis in L7-PP2B KO mice across various ages, we demonstrate that, in contrast to early postnatal loss of function, acute pharmacological blockage of PP2B in adult mice affected the adaptation of the vestibulo-ocular reflex (VOR), but not the reflex itself. Moreover, we found that adult-onset, but prolonged deletion of PP2B in adult mice, using a tamoxifen-dependent inducible model, affects all forms of plasticity and the OKR, but again not the VOR itself. In contrast, juvenile L7-PP2B KO mice exhibit a progressive loss of PP2B and concurrent changes in the VOR, in addition to the loss of adaptive abilities. Together, our results suggest that adult-onset loss of plasticity affects adaptation, whereas a loss during development results in an alternative set-point of the basal reflex.

## Materials and Methods

All experiments were approved by the Dutch Ethical Committee for animal experiments and were in accordance with the Institutional Animal Care and Use Committee.

### Animals

For all experiments, we used adult male and female mice with a C57Bl/6 background that were, unless stated otherwise, group housed until the first surgery and then individually housed. All mice had food ab libitum and were on a 12:12 light/dark cycle. In all experiments the experimenters were blind to mouse genotypes. Mice with PC-specific knockout of PP2B were used previously (Lin Z, et al., 2021; Schonewille M, et al., 2010). In short, the regulatory subunit of the calcium/calmodulin-activated protein phosphatase 2B was flanked with loxP sites (MGI: C57BL/6-Ppp3r1^tm1Stl^/J, also referred to as Ppp3r1^fl/fl^), which can be selectively deleted from Purkinje cells by crossing this line with the pcp2-promotor driven Cre recombinase expression (MGI: Tg(Pcp2-cre)1Amc/J, also known as L7^cre^). Mice of the following genotypes were used for the experiments: L7-cre^+/-^/PPP3R1^fl/fl^ (referred to as L7-PP2B KO) and L7-cre^-/-^/PPP3R1^fl/fl^, L7-cre^+/-^/PPP3R1^+/+^ and L7-cre^+/-^/PPP3R1^+/+^ (all littermate controls or L7-PP2B Ctrl). Eye movement recordings were performed in adult mice aged between 10-30 weeks of age, and in two juvenile cohorts: aged postnatal 18-21 days (P18-21) and aged postnatal 26-30 days (P26-30). Inducible PC-specific PP2B knockouts were generated by crossbreeding mice carrying the floxed PPP3R1 with mice expressing the tamoxifen-sensitive Cre recombinase Cre-ERT2 under the control of the L7 promoter (MGI: Tg(Pcp2-cre/ERT2)17.8Ics), generating experimental mice: L7^Cre-ERT2/-^; PPP3R1^fl/fl^, referred to as iL7-PP2B cKO + TAM. Tamoxifen was dissolved in corn oil to obtain a 20 mg/ml solution, and intraperitoneally injected into all mice, mutants and controls, for consecutive 5 days, four weeks prior to eye movement recordings. Injections were performed in adults between 24-40 weeks of age. Experimental cohorts were always injected at the same time during the day. Littermate controls were mice without L7^Cre-ERT2^ expression, but with tamoxifen injections (L7^-/-^; PPP3R1^fl/fl^, referred to as iL7-PP2B Ctrl + Tam) or mice with L7^Cre-ERT2^, but without vehicle instead of tamoxifen injections (L7^Cre-ERT2+/-^; PPP3R1^fl/fl^, referred to as iL7-PP2B cKO + Veh).

### Immunohistochemistry

Anesthetized mice were perfused with 4% paraformaldehyde in 0.12M phosphate buffer (PB). Brains were taken out and post-fixed for 1 h in 4% PFA at room temperature, then transferred in 10% sucrose overnight at 4° C. The next day, the solution was changed for 30% sucrose and left overnight at 4° C. Non-embedded brains were sectioned either sagittally or transversally at 40µ m thickness with freezing microtome. Free-floating sections were rinsed with 0.1M PB and incubated 2h in 10mM sodium citrate at 80° C for 2 h, for antigen retrieval. For immuno-fluorescence, sections were rinsed with 0.1M PB, followed by 30 minutes in Phosphate Buffered saline (PBS). Sections were incubated 90 minutes at room temperature in a solution of PBS/0.5% Triton-X100/10% normal horse serum to block nonspecific protein-binding sites, and incubated 48 h at 4° C in a solution of PBS/0.4% Triton-X100/2% normal horse serum, with primary antibodies as follows: Aldolase C (1:500, goat polyclona, SC-12065), Calbindin (1:7000, mouse monoclonal, Sigma, # C9848), and PP2B (1:500, rabbit polyclonal, proteintech). After rinsing in PBS, sections were incubated 2 h at room temperature in PBS/0.4% Triton-X100/2% normal horse serum solution with secondary antibodies coupled with Alexa488, Cy3 or Cy5 (Jackson ImmunoResearch), at a concentration of 1:200. Sections were mounted on coverslip in chrome alum (gelatin/chromate) and covered with Mowiol (Polysciences Inc.). Images were acquired with an upright LSM 700 confocal microscope (Zeiss).

### Western blot

Cerebellar tissue from mice was dissected and immediately frozen in liquid nitrogen. Samples were homogenized with a Dounce homogenizer in lysis buffer containing 50 mM Tris-HCl pH 8, 150 mM NaCl, 1% Triton X-100, 0.5% sodium deoxycholate, 0.1% SDS and protease inhibitor cocktail. Protein concentrations were measured using Pierce BCA protein assay kit (Thermo Fisher). Samples were denatured and proteins were separated by SDS-PAGE in Criterion™ TGX Stain-Free™ Gels (Bio-Rad), and transferred onto nitrocellulose membranes with the Trans-Blot® Turbo™ Blotting System (Bio-Rad). Membranes were blocked with 5% BSA (Sigma-Aldrich) in TBST (20mM Tris-HCl pH7.5, 150mM NaCl and 0.1%, Tween20) for 1 h and probed with the primary antibodies as follows: Dyn1-S778 (1:1000, sheep monoclonal, Abcam), Dyn1 (1:1000, mouse monoclonal, Santa cruz), and probed with secondary antibodies subsequently. Membranes were scanned by Odyssey Imager (LI-COR Biosciences) and quantified using Image Studio Lite (LI-COR Biosciences).

### Stereotaxic AAV-injections

To re-express PP2B we performed stereotaxic injection of the AAVs containing floxed, cre-dependent native PP2B (AAV-CAG-eGFP-PP2B) or enzyme-dead H151A mutant PP2B constructs (AAV-CAG-eGFP-PP2B/H151A) two weeks prior to eye movement measurements in the bilateral flocculi of adult L7-PP2B KO mice following a lateral-dorso-ventral direct approach (1 mm depth; 100-150 nl, titres 1.0-1.2 × 10^13^ vg ml^-1^). After injection, the pipette was left in place for > 10 min before being slowly withdraw. All the process was done under isoflurane (4% induction, 1.5-2% maintenance) anesthesia. Only mice with at least 30% but typically ≥ 50% of the flocculus covered by the injection were included in the results.

### Compensatory eye movement recordings

Mice were subjected to compensatory eye movement recordings as previously described in detail (Schonewille M, et al., 2010). In short, mice were equipped with an immobilizing construct, or ‘ pedestal’, under general anesthesia with isoflurane/O_2_. After 2-3 days of recovery, mice were head-fixed with the body loosely restrained in a custom-made restrainer and placed in the center of a turntable (diameter: 60 cm) in the experimental set-up (**Fig. 1**). A round screen (diameter 63 cm) with a random dotted pattern (‘ drum’) surrounded the mouse during the experiment. Compensatory eye movements were induced by sinusoidal rotation of the drum in light (OKR), rotation of the table in the dark (VOR) or the rotation of the table in the light (visually enhanced VOR, VVOR) with an amplitude of 5° at 0.1-1 Hz. Motor performance in response to these stimulations was evaluated by fitting the averaged stimulus and responses with a sine and calculating the gain (fitted amplitude eye velocity/stimulus velocity) and phase (timing of eye relative to stimulus in degrees). Motor learning was studied by subjecting mice to mismatched visual and vestibular input. Rotating the drum (visual) and table (vestibular) simultaneously, in-phase at 0.6 Hz (both with an amplitude of 5°, 5 x 10 min) in the light will induce a decrease of the gain of the VOR (in the dark). Subsequently, VOR Phase reversal was tested by continuing the next days (day 2-5, keeping mice in the dark in between experiments) with in-phase stimulation, but now with drum amplitudes of 7.5° (days 2) and 10° (days 3, 4, and 5), while the amplitude of the turntable remained 5°. This resulted, over days of training, in the reversal of the VOR direction, from a normal compensatory rightward eye movement (in the dark), when the head turns left, to a reversed response with a leftward eye movement, when the head moves left. At the end of the VOR phase reversal training the OKR was probed again and compared to the OKR before training, to examine OKR gain increase. VOR gain increase was evoked by subjecting mice to out of phase drum and table stimulation at 1.0 Hz (both with an amplitude of 1.6°). A CCD camera was fixed to the turntable in order to monitor the eyes of the mice. Eye movements were recorded with eye-tracking software (ETL-200, ISCAN systems, Burlington, NA, USA). For eye illumination during the experiments, two infrared emitters (output 600LJmW, dispersion angle 7°, peak wavelength 880 nm) were fixed to the table and a third emitter, which produced the tracked corneal reflection as a reference point, was mounted to the camera and aligned horizontally with the optical axis of the camera. Eye movements were calibrated by moving the camera left-right (peak-to-peak 20°) during periods that the eye did not move. Gain and phase values of eye movements were calculated using custom-made Matlab routines (MathWorks, https://github.com/MSchonewille/iMove).

**Figure 1:**
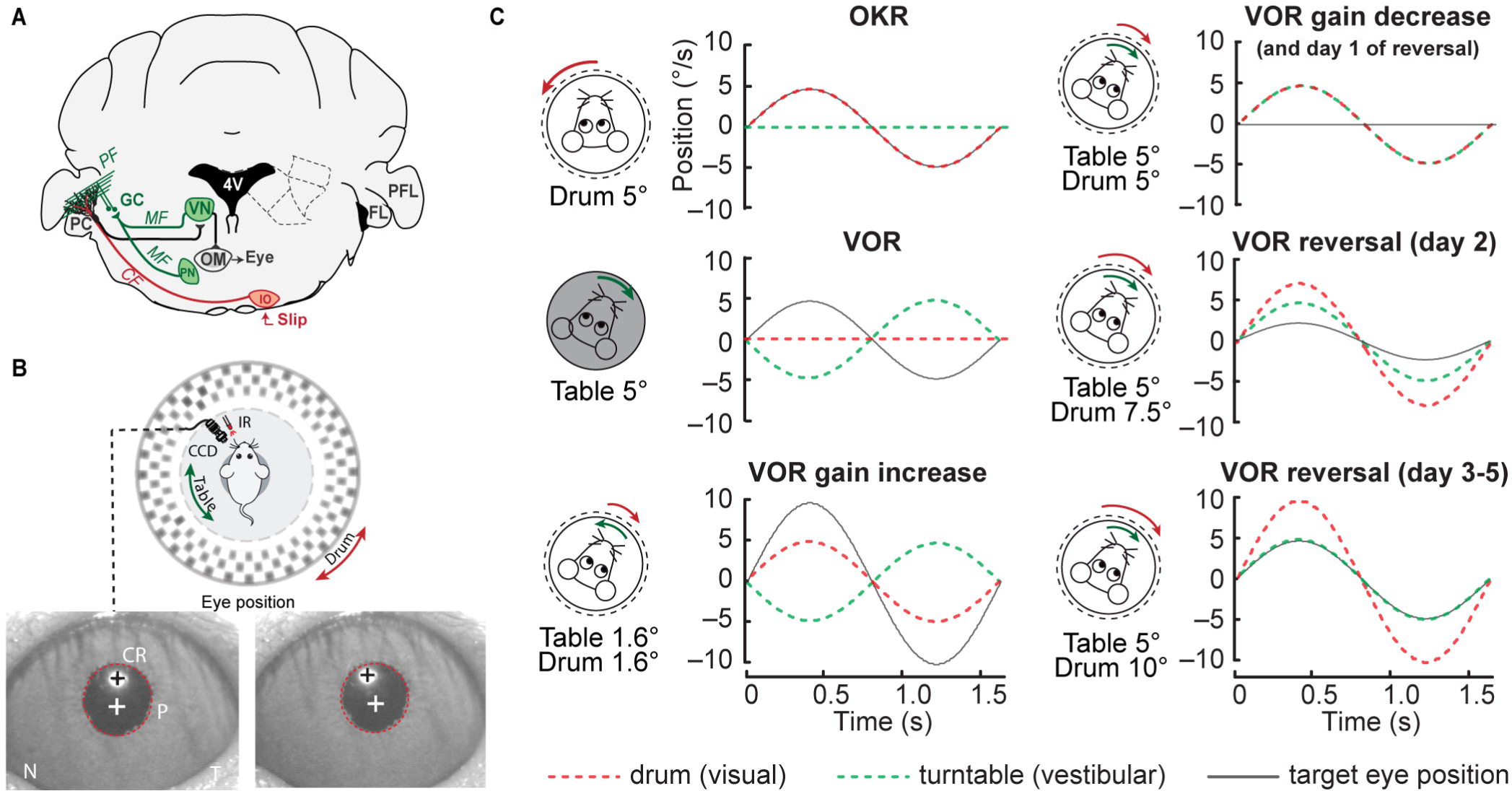
Schematic illustration of the methodology for recording compensatory eye movement and its adaptation. **A**, Cerebellar circuitry controlling compensatory eye movements and their adaptation. PCs in the flocculus (FL) receive vestibular and visual input via the mossy fiber (MF) - parallel fiber (PF) system (green) and climbing fiber input (CF, red) from the inferior olive (IO), indicating retinal slip. By integrating these two inputs, PCs adjust eye movements via the vestibular nuclei (VN) and the oculomotor (OM) neurons. PN, pontine nuclei; GC, granule cell. **B,** Eye movement recording setup. Mice are head-fixed in the center of a turntable for vestibular stimulation and surrounded by a random dotted pattern (‘ drum’) for visual stimulation. A CCD camera was used for infrared (IR) video-tracking of the left eye. Bottom, examples of nasal (N) and temporal (T) eye positions. Red circles, pupil fit; black cross, corneal reflection (CR); white cross, pupil center. **C**, Graphical explanation of the trajectory relationship between the turntable (equivalent to vestibular stimulation, green dashed line), drum (equivalent to visual stimulation, red dashed line) and target eye trace (black line) for basal compensatory eye movements (OKR and VOR) and VOR adaption paradigms (gain-increase, gain-decrease and phase reversal).

### Statistics

All values are shown as mean ± SEM. As described previously (Wu B et al., 2019), the behavioral experiments group sizes were estimated a priori using sample size calculations based on minimal relevant differences and expected variation in control cells or mice. For compensatory eye movements across stimulation frequencies and adaptation over time, an ANOVA for repeated measures was used to determine statistical significance, followed by posthoc tests for individual comparisons. For the complete dataset, see **Extended Data**: **Table 1-6**. All statistical analyses were performed using SPSS 20.0 software. Data was considered statistically significant if P < 0.05. (* indicates *P* < LJ0.05, ** indicates *P* < LJ0.01, *** indicates *P*LJ< LJ0.001).

## Results

Previous work has shown that chronic genetic ablation of native PP2B from Purkinje cells (PCs) in L7-PP2B KO mice affects both the performance of compensatory eye movement reflexes as well as the plasticity-dependent adaptation thereof (Schonewille M, et al., 2010), (Figure 1). To disentangled the contribution of the acute absence of the gene during the experiment from the long-term effects of genetic deletion, we first aimed to acutely block PP2B. Recent work indicates that PP2B facilitates the motor learning predominantly through its enzymatic activity (Lin Z, et al., 2021). Therefore, we first set out to test the effects of acute disruption of inhibiting the enzymatic activity by pharmacological intervention.

### Acute pharmacological blocking of PP2B partially affects the adaptation of VOR but not the reflex itself

To test the effect of acutely blocking the enzymatic activity of PP2B, we administered FK506, a selective PP2B inhibitor (Butcher SP et al., 1997; Pardo R et al., 2006). L7-PP2B control littermate mice (L7-PP2B Ctrl) were injected intraperitoneally 15 min before start of the experiment with 10 mg/kg FK506 or vehicle (10% DMSO, 10% ethanol in 0.9% saline). For comparison L7-PP2B mutant mice (L7-PP2B KO) were also injected with vehicle solution **(Fig. 2A)**. In line with previous studies (Clayton EL et al., 2009; Cottrell JR et al., 2013), the pharmacological inhibition of PP2B by FK506 resulted in a similar hyper-phosphorylation of Ser778 in Dynamin1, a presynaptic GTPase dephosphorylated by PP2B, to that of genetic deletion of PP2B, indicating a comparable blockage of the enzymatic activity of PP2B **(Fig. 2B)**. L7-PP2B KO mice injected with vehicle exhibited significant deficits in the gain of the optokinetic reflex (OKR, p=0.042, red vs black, repeated measure ANOVA with Bonferroni correction) and correlating increase in the vestibulo-ocular reflex gain (VOR gain, p=0.013, red vs green) across different frequencies, while injecting control mice with FK506 did not significantly affect motor performance in either OKR or VOR gain (both p=1.000, green vs black, **Fig. 2C-D**). While wildtype mice injected with FK506 and mice injected with vehicle learned equally well in gain-decrease paradigm (p=1.000, green vs black, **Fig. 2E**), FK506 caused a significant impairment in the phase change induced by VOR phase-reversal training over 5 consecutive days (p<0.001, green vs black, **Fig. 2F**). In line with previous results, L7-PP2B KO mutant mice injected with vehicle showed learning deficits in both paradigms (all p<0.01, red vs black or red vs green, **Fig. 2E-F**). Thus, acute pharmacological blocking of the enzymatic function of PP2B only partially affects VOR adaptation, while leaving compensatory eye reflexes intact.

**Figure 2:**
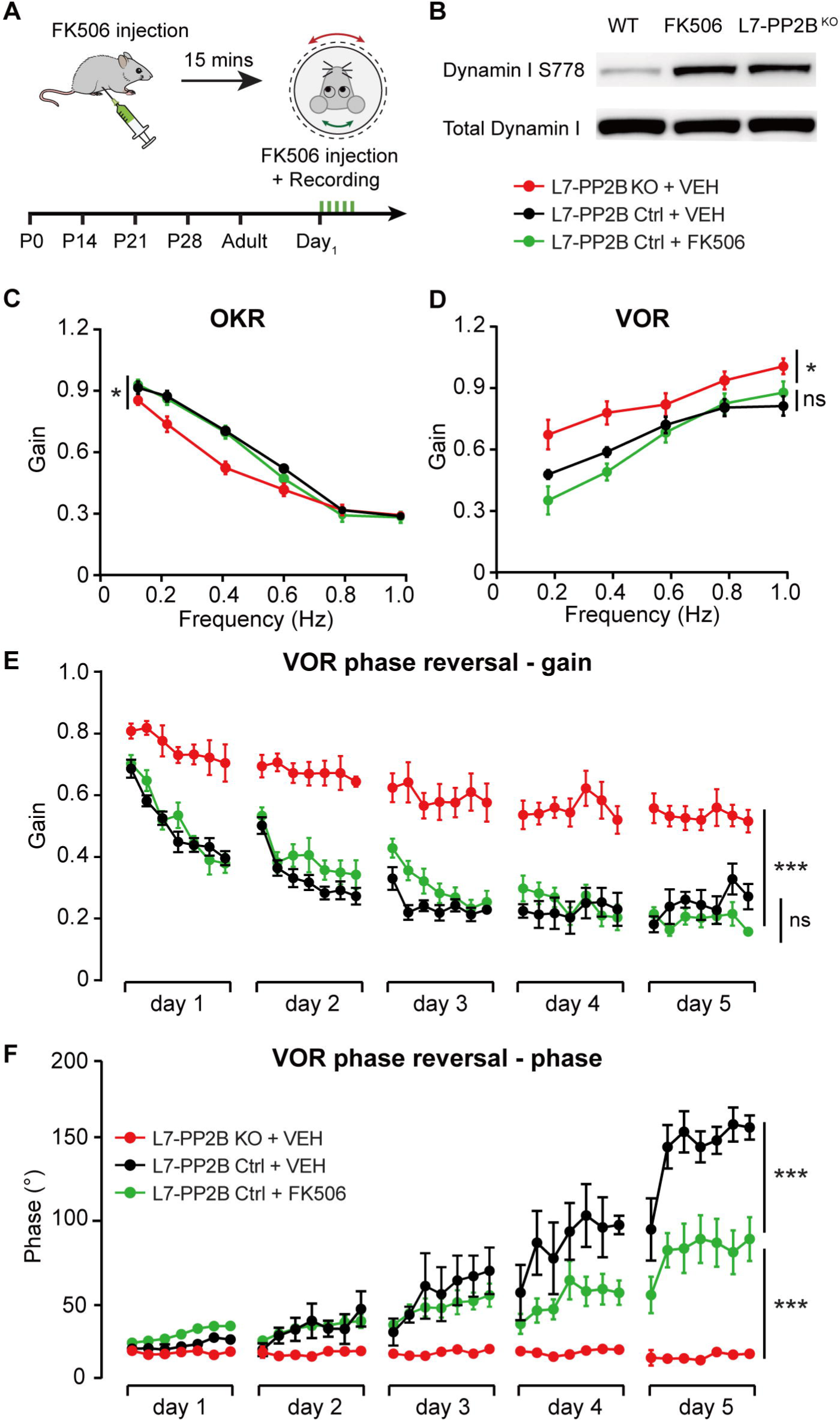
Effects of acute pharmacological blocking of PP2B on reflex and reflex adaptation. **A**, Schematic strategy of eye movement recording following i.p. FK506 injection. **B**, Western blot of Ser778 in Dynamin1 validating FK506 inhibition efficiency. Cortices tissue was collected from mice after the behavioral experiments. **C-F**, Recorded OKR gain (**C**), VOR gain (**D**), VOR gain decrease learning (**E**) and VOR phase reversal training across consecutive 5 days (**F**) in L7-PP2B KO wildtype littermate mice injected with FK506 (green, n=8 mice) and vehicle (black, n=7 mice), and L7-PP2B KO mutant mice injected with vehicle (red, n=6 mice), respectively. Panels **E** and **F** are re-used from Lin Z et al., 2021 (Figure 1C), but re-plotted in gain and phase plots to facilitate comparison with later figures. Data are represented as mean ± SEM. ns, non-significant; \**p* < 0.05; \*\**p* < 0.01; \*\*\**p* < 0.001 (see Extended Data Table 2-1 for details).

### Deletion of PP2B in adulthood affects OKR performance and VOR adaptation, but not basal VOR

As acute interventions were not able to replicate the phenotype of early postnatal loss of PP2B, we next asked if delayed genetic PP2B deletion, starting from adulthood instead of early development, could affect the behavior of compensatory eye movements. To test this, we crossed the *loxP*-flanked PP2B mice with tamoxifen-dependent L7^Cre-ERT2^ to generate conditional iL7-PP2B cKO mice (**Fig. 3A**). When mice were more than 3 months old, we injected iL7-PP2B cKO mutant mice and iL7-PP2B Ctrl control mice with tamoxifen, as well as iL7-PP2B cKO mice with vehicle alone and waited for >4 weeks before starting experiments. If the deficits of eye movement baseline reflexes and plasticity were completely or in part of developmental origin, we should observe no or less changes in adult conditional L7-PP2B KO mice after tamoxifen injections. Delayed genetic deletion of PP2B had profound effects on performance and learning. First, after tamoxifen injection OKR gain was impaired in iL7-PP2B cKO + TAM mice compared to iL7-PP2B Ctrl + TAM mice (**Fig. 3B,** p=0.020). Similarly, motor learning affected eye movement adaptation in gain-increase (p=0.047), gain-decrease (p=0.001) and VOR phase reversal (day 4, p<0.001) (**Fig. 3D-F**, black vs red). In contrast to OKR and VOR adaptation, however, the VOR gain of tamoxifen injected iL7-PP2B cKO + TAM mice was not affected (**Fig. 3C**, p=0.375), the only deficit not replicated after adult-onset deletion. As expected, iL7-PP2B cKO + TAM mice did not differ from vehicle injected mice in any of the OKR and VOR (adaptation) tests (**Fig. 3B** and **3D-F**, blue vs black). Finally, we confirmed that 4 weeks after tamoxifen treatment PP2B is ablated from PCs in iL7-PP2B cKO + TAM mice (**Fig. 3G**), compared to iL7-PP2B Ctrl + TAM mice injected with tamoxifen (**Fig. 3H**) or iL7-PP2B cKO + VEH mice injected with vehicle (**Fig. 3I**).

**Figure 3:**
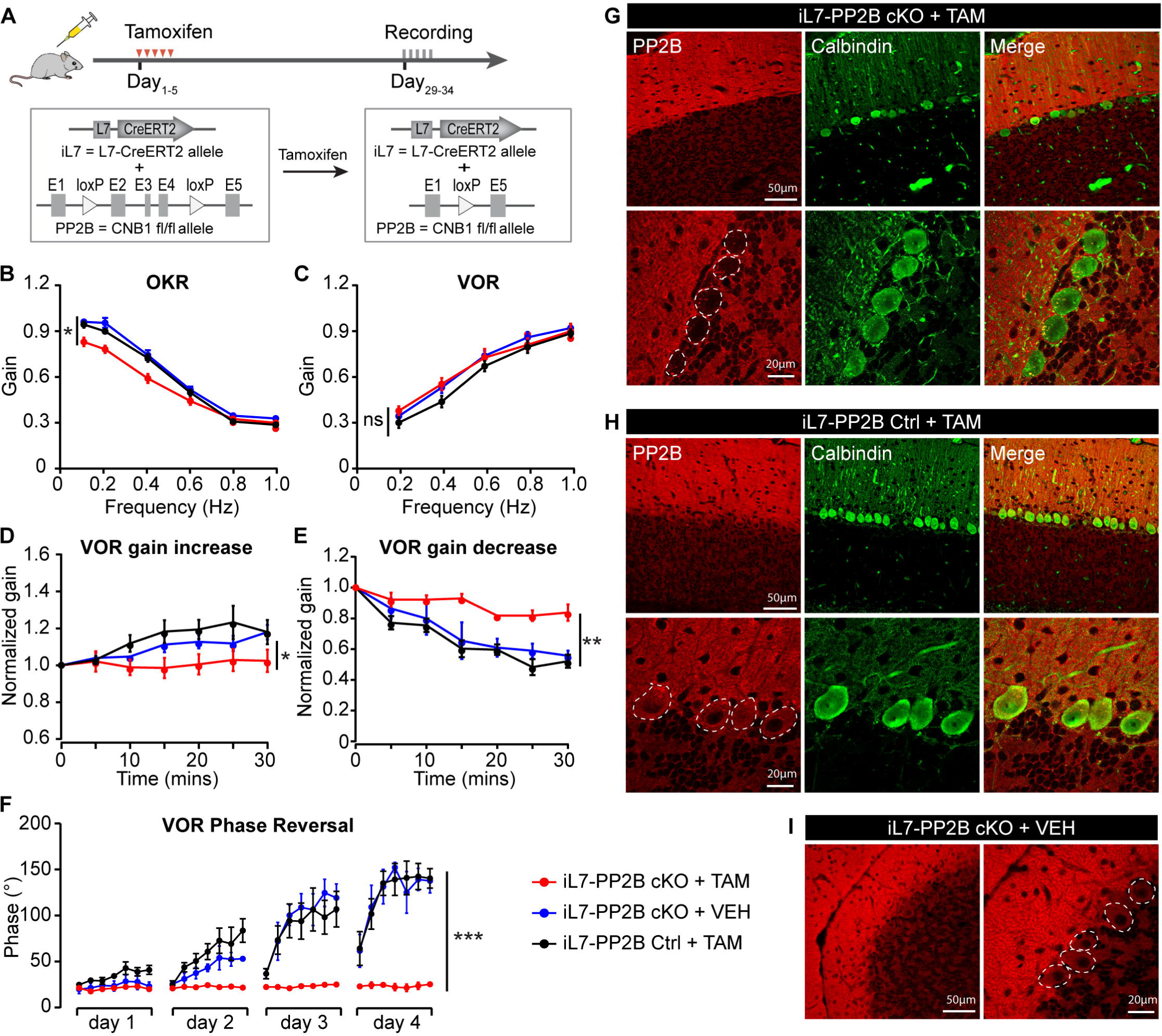
Effects of Purkinje cell-specific adult-onset genetic ablation of PP2B. **A**, Schematic strategy for generation of a PC-specific PP2B deletion triggered by tamoxifen. **B-F**, Recorded OKR gain (**B**), VOR gain (**C**), VOR gain increase learning (**D**), VOR gain decrease learning (**E**) and VOR phase reversal training across consecutive 4 days (**F**) in iL7-PP2B cKO + TAM mice (red, n=12), iL7-PP2B Ctrl + TAM mice (black, n=13) injected with tamoxifen and iL7-PP2B cKO + VEH mice injected with vehicle-only (blue, n=10). Dotted red lines indicate the results previously obtained in L7-PP2B KO mice for reference. **G-I**, Corresponding immunofluorescent images of the three experimental groups (red, PP2B; green, calbindin; yellow, merge). White dotted lines indicate the somata of Purkinje cells. Data are represented as mean ± SEM. ns, non-significant; \**p* < 0.05; \*\**p* < 0.01; \*\*\**p* < 0.001 (see Extended Data Table 3-1 for details).

Taken together, we find that prolonged PP2B deletion after cerebellar maturation is able to reproduce the deficits observed in OKR and VOR adaptation following developmental deletion of PP2B in L7-PP2B KO mice (red dotted line, **Fig. 3C**), but not the deficit in VOR gain.

### PP2B ablation affects eye movement behavior during early development

To investigate the behavioral consequences in relation to the onset and temporal profile of PP2B deletion, we examined eye movements in juvenile L7-PP2B KO mice during the first four postnatal weeks. Given that the L7-Cre promotor starts to be expressed from the first postnatal week, with a complete expression in all PCs by the end of postnatal week 2 (Barski JJ et al., 2000), we examined the expression level of the PP2B protein in the flocculus of L7-PP2B KO mice on various days at the juvenile age of postnatal (P) 14, 18, 20 and 24 days. As early as the age of P14, when mice started to open their eyes, a small proportion (∼20%) of floccular PCs no longer expressed PP2B (**Fig. 4A**). At P18, the proportion of PCs with deleted PP2B became larger (∼50%, **Fig. 4B**). From P20 towards P24, over 90% of PCs lacked PP2B, approaching the virtually complete deletion observed in adult L7-PP2B KO mice (**Fig. 4C-D**).

**Figure 4:**
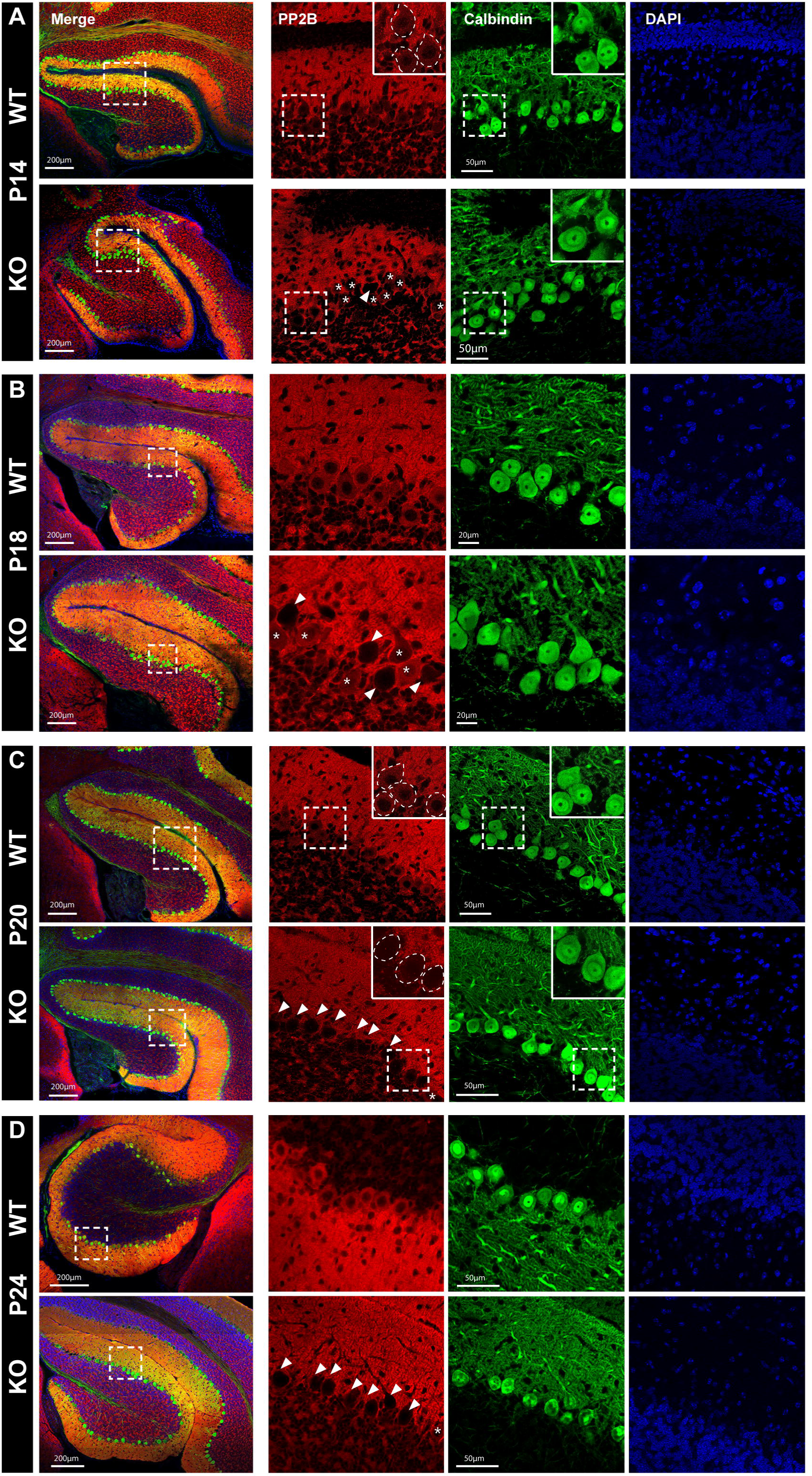
Floccular PP2B expression across development in juvenile L7-PP2B KO mice. **A-D**, Immunofluorescent images of PP2B expression in the flocculus of L7-PP2B KO mice at postnatal day (P) 14 (**A**), 18 (**B**), 20 (**C**) and 24 (**D**). Leftmost column, overview staining of flocculi; boxed areas are magnified in the three columns on the right (red, PP2B; green, calbindin; blue, DAPI). White dotted lines indicate the somata of PCs. White triangles, PC somata without PP2B; white asterisks, PC somata with PP2B. Scale bars indicate 200 μ m for merge images, 50 μ m for other images in A, C and D and 20 μ m for those in B.

Next, we tested compensatory eye movement in juvenile L7-PP2B KO and control mice at P18-21, P26-30 and compared the results to adult mice. At P18-21, L7-PP2B KO mice showed intact baseline performance of OKR and VOR, as well as normal VOR gain increase and gain decrease learning, compared with their littermate controls (OKR, p=0.965; VOR, p=0.461; VOR gain increase, p=0.636; VOR gain decrease, p=0.106, squared lines, **Fig. 5A-D**). Note that in our histological analysis we found that ∼50% of the floccular PCs PP2B was deleted at this stage, yet there is no evidence for an effect on the compensatory eye movement adaptation, possibly indicating that there is sufficient remaining circuit plasticity despite a substantial reduction of PP2B. In contrast, at P26-30 L7-PP2B KO mice showed evident impairments not only in OKR (p=0.001) and VOR (p=0.013) performance (triangular lines, **Fig. 5A-B**), but also in VOR gain increase (p=0.014) and VOR gain decrease (p=0.026) learning as well as VOR phase reversal (p<0.001) (triangular lines, **Fig. 5C-E**). Behavior at P26-P30 is similar to that observed in adult L7-PP2B KO mice (data from Schonewille et al., 2010; here shown in dotted lines, **Fig. 5A-E**) with the most prominent difference being that learning rates are lower in adult mice (Beekhof GC, et al., 2021). Note that some capacity for learning was still observed in the juvenile L7-PP2B KO mice aged P26-30 (red triangular line, **Fig. 5E**). Together, these findings indicate a direct correlation between the absence of PP2B and behavioral deficits. Moreover, at four weeks of age there is a virtually complete deletion of PP2B deletion that affects all recorded reflexes and adaptation, including the VOR, indicating that the size of this innate reflex is set early in development.

**Figure 5:**
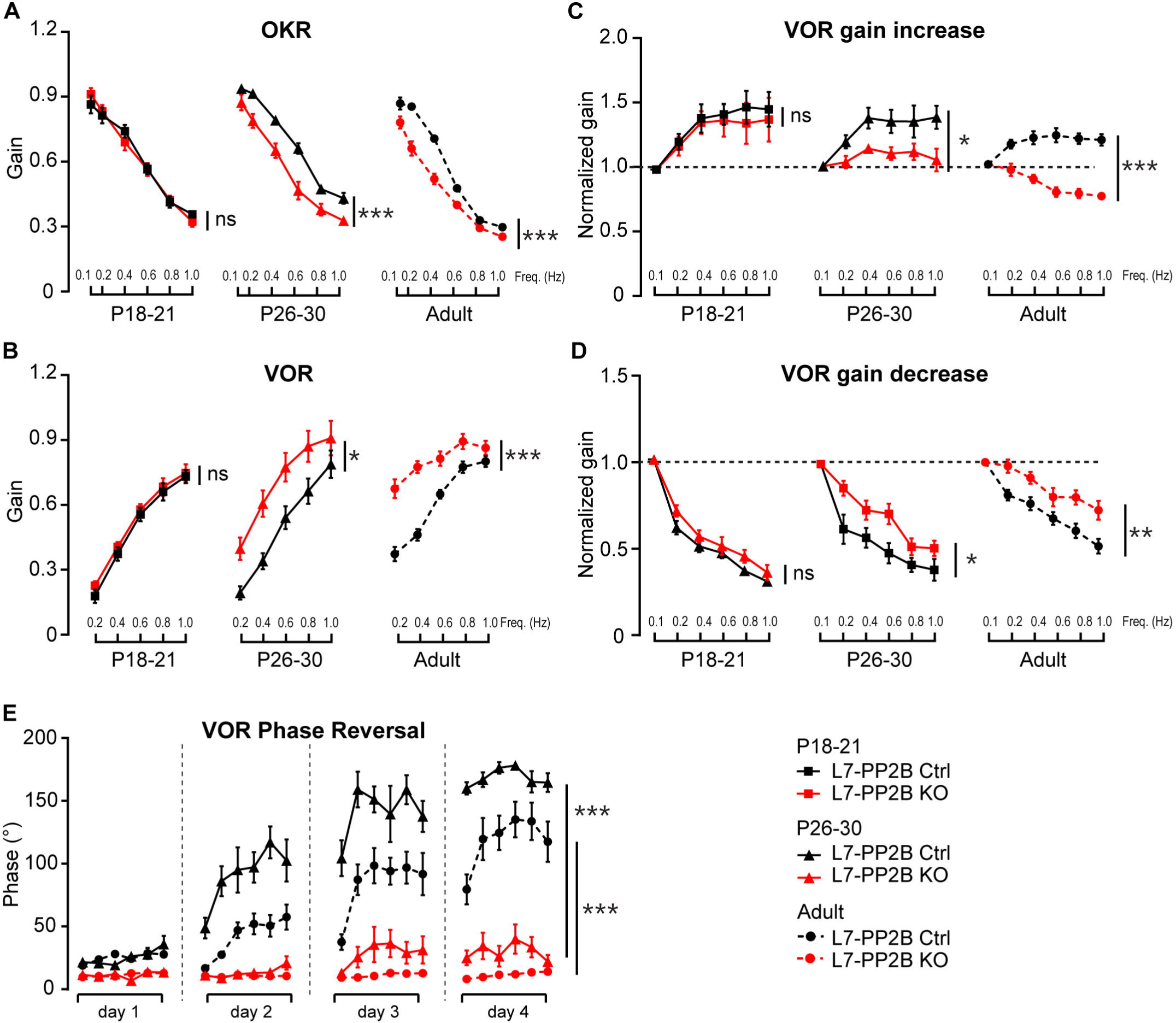
Comparison of reflex and reflex adaptation in L7-PP2B KO mice at different developmental age. **A**, OKR gain in L7-PP2B KO and their littermate control mice at the age of P18-21 (KO, n=9 mice; WT, n=11 mice; squared lines), P26-30 (KO, n=10 mice; WT, n=13 mice; triangular lines) and adulthood (KO, n=15 mice; WT, n=23 mice; dotted cycled lines, data from Schonewille et al., 2010). **B**, same as **A**, but for VOR gain in mice aged P18-21(KO, n=8 mice; WT, n=11 mice; squared lines), P26-30 (KO, n=6 mice; WT, n=7 mice; triangular lines) and adulthood (KO, n=14 mice; WT, n=23 mice; dotted cycled lines). **C**, VOR gain increase adaptation in L7-PP2B KO and their littermate control mice at the age of P18-21(KO, n=9 mice; WT, n=10 mice; squared lines), P26-30 (KO, n=9 mice; WT, n=7 mice; triangular lines) and adulthood (KO, n=10 mice; WT, n=13 mice; dotted cycled lines). **D**, same as **C**, but for VOR gain decrease adaptation in mice aged P18-21(KO, n=8 mice; WT, n=7 mice; squared lines), P26-30 (KO, n=7 mice; WT, n=8 mice; triangular lines) and adulthood (KO, n=13 mice; WT, n=13 mice; dotted cycled lines). **E**, Phase-reversal training in mice aged P26-30 (KO, n=10 mice; WT, n=11 mice; triangular lines) and adulthood (KO, n=8 mice; WT, n=8 mice; dotted cycled lines). Littermate controls in black; mutant mice in red; Data are represented as mean ± SEM. ns, non-significant; \**p* < 0.05; \*\**p* < 0.01; \*\*\**p* < 0.001 (see Extended Data Table 5-1, 5-2, 5-3 for details).

### Restoring PP2B expression in adult L7-PP2B KO mice rescues VOR adaptation and OKR, but not VOR

Finally, if adult-onset deletion of PP2B will prevent VOR gain from being affected, the inverse should also hold: juvenile deletion followed by re-expressing PP2B in adult mice should rescue adaptation, while VOR gain remains impaired. To test this hypothesis, we injected restored PP2B expression in L7-PP2B KO mice by injecting AAV1-CAG-eGFP-PP2B and tested baseline compensatory eye movements and VOR adaptation. For comparison, in another group of L7-PP2B KO mice with AAV1-CAG-eGFP-PP2B/H151A, which lacks its enzymatic function due to the H151A mutation (Baksh S et al., 2000; Lin Z, et al., 2021). AAV injections were made bilaterally in the flocculi and the location of injection was confirmed by post-mortem analysis (**Fig. 6A-B**). Re-expression of functional PP2B in L7-PP2B KO mice increased the OKR gain compared to that of PP2B-H151A, resulting in levels that are comparable to controls (p=0.997, purple vs black, repeated measure ANOVA with Bonferroni correction, **Fig. 6C**). Although in VOR phase reversal the improvement in phase was not significant (eg. p=0.426 for day 5, purple vs red, **Fig. 6E**) the ability to decrease the gain, as a first step in the reversal, is significantly improved (all p<0.05 for day 1-5, purple vs red, **Fig. 6F**). The expression of PP2B lacking its enzymatic function had virtually no effect on OKR gain, VOR gain or VOR adaptation (compared with red curves, **Fig. 2C-F**) (Lin Z, et al., 2021). Unlike the effect on VOR adaptation and OKR gain and in line with our hypothesis, the re-expression of fully functional PP2B in L7-PP2B KO mice did not significantly change the VOR gain. As a result, VOR gain differs from that in L7-PP2B Ctrl (p=0.028, purple vs black, **Fig. 6D**) and not from L7-PP2B KO mice injected with PP2B-H151A (p=0.929, purple vs red, **Fig. 6D**).

**Figure 6:**
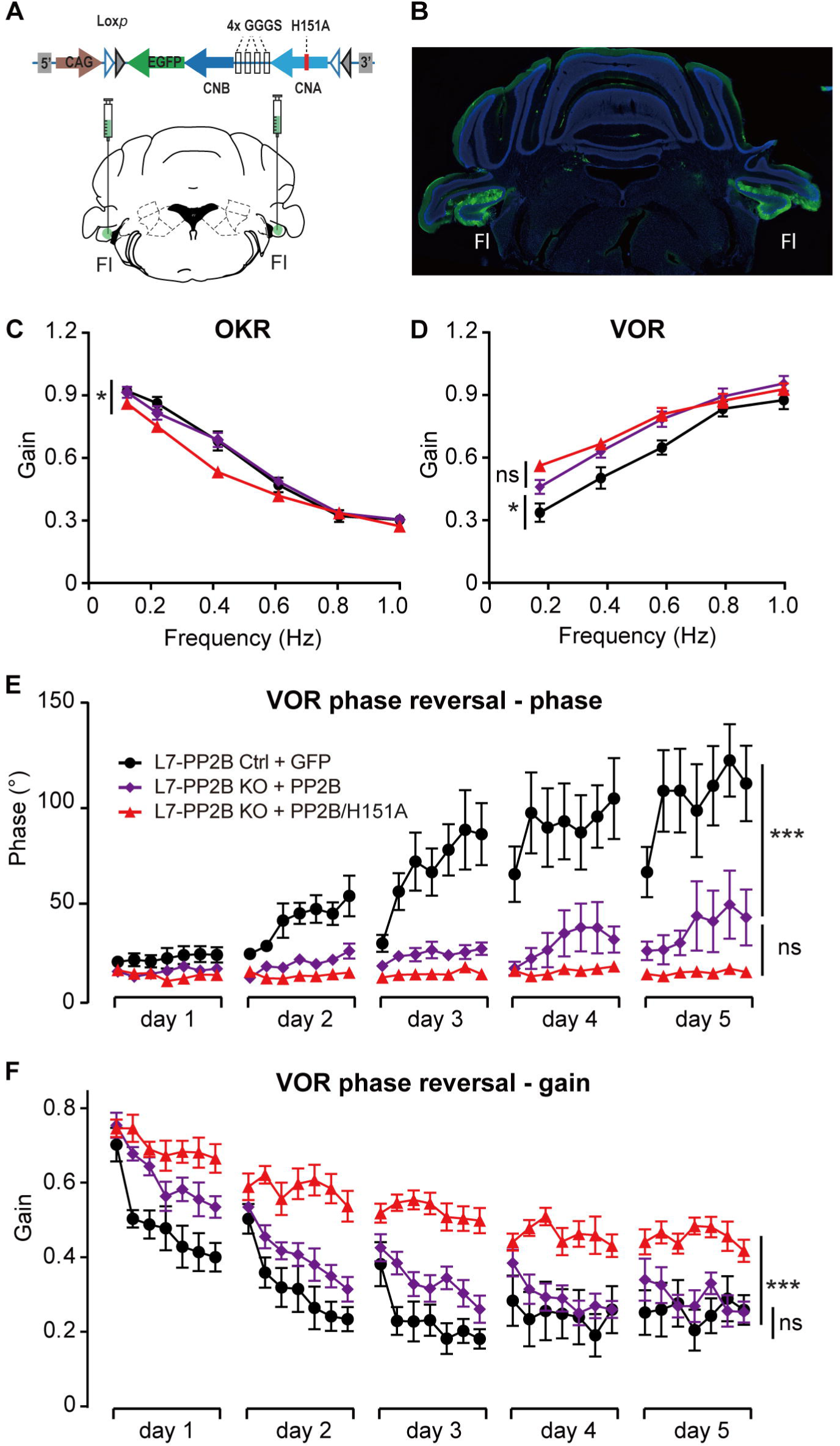
Re-expressing functional PP2B and PP2B lacking enzymatic function in adult L7-PP2B KO mice. **A**, Schematic depiction of the AAVs carrying Cre-dependent functional PP2B and enzyme-dead PP2B due to the H151A mutation (PP2B/H151A), and the strategy of AAV injections made bilaterally in the flocculi. **B**, Example of AAVs injections with either functional PP2B or enzyme-dead PP2B/H151A in bilateral flocculi (visualized by the co-expression of EGFP). **C-F**, Eye movement recording of OKR gain (**C**), VOR gain (**D**), VOR phase reversal with the change in phase (**E**) and gain (**F**) across consecutive 5 days in L7-PP2B control mice injected with AAVs with GFP (black, n=9 mice) and L7-PP2B KO mutant mice injected with AAVs with PP2B (purple, n=11 mice) or PP2B/H151A (red, n=12 mice), respectively. Data are represented as mean ± SEM. ns, non-significant; \**p* < 0.05; \*\**p* < 0.01; \*\*\**p* < 0.001 (see Extended Data Table 6-1 for details).

Taken together, we find that re-instating PP2B in the majority of floccular Purkinje cells of adult L7-PP2B KO mice is sufficient to rescue OKR gain and improve VOR adaptation, but does not significantly affect the VOR gain. These results confirm that the setpoint of the vestibular-input driven innate reflex is largely, if not completely, set in the first weeks after eye opening.

## Discussion

The present study probed the contribution of the cerebellar cortex, in particular Purkinje cell plasticity, to the formation and maintenance of a reflexive behavior. We found that an acute block of PP2B function, an enzyme required for synaptic and non-synaptic plasticity, did not affect the baseline reflexes of compensatory eye movements, which is in clear contrast with what happens after permanent genetic deletion at early development. Prolonged Purkinje cell - specific genetic PP2B ablation, starting from the adult stage, selectively affected VOR adaptation and basal OKR, but not basal VOR, performance. In contrast, at the age of P26-30, shortly after the complete loss of PP2B from their Purkinje cells, juvenile L7-PP2B KO mice exhibited impairments in OKR, VOR and VOR phase reversal comparable with their adult mutant counterparts. Finally, re-expressing PP2B in adult L7-PP2B KO mice rescued OKR gain and VOR adaptation, but did not rescue the VOR gain (see summary in **Fig. 7**). Thus, using temporal variations in PP2B ablations, we demonstrate that disruption or absence of PP2B always directly impairs adaptation, while the setpoint of the VOR is exclusively determined during cerebellar development in the juvenile stage.

**Figure 7:**
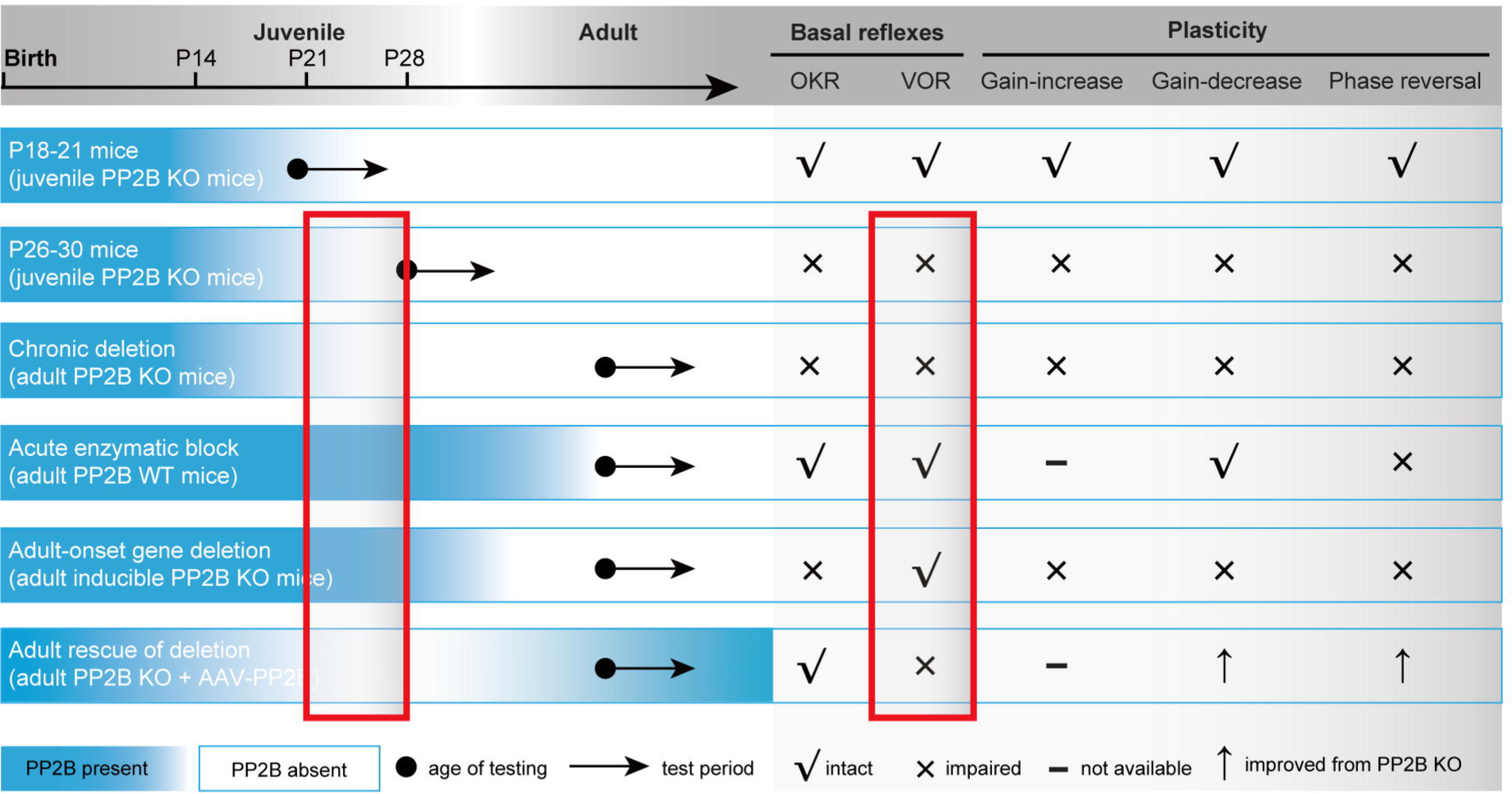
Summary of various impacts of PP2B manipulation strategies on both basal reflexes and plasticity of compensatory eye movement in juvenile or adult mouse. Note that all behavioral tests are normal in juvenile mice at P18-21. However, at P26-30 juvenile PP2B KO mice showed evident deficits in all basal reflexes and plasticity measurements, similar to adult PP2B KO mice. Conversely, when PP2B was acutely or chronically blocked, but with adult-onset, all reflexes and reflex adaptations, except for the VOR gain were affected. Moreover, re-expression functional PP2B improved OKR gain and elements of VOR adaptation, but not VOR gain.

Compensatory eye movements form the ideal substrate to study the ontogenetic steps related to the onset and maturation of reflexes as well as their adaptations. Two processes together assure adequate compensatory eye movements throughout life: 1) the reflexes and related circuit need to be configurated during development, and 2) the adaptive, plastic process needs to maintain optimal performance by continuously calibrating the response to reduce retinal slip during head perturbations (Alahyane N et al., 2016; Charpiot A et al., 2010; Finocchio DV et al., 1991; Roucoux A et al., 1983). The flocculus of the cerebellum is required for proper setup and adaptation of the vestibulo-ocular reflex (Charpiot A, et al., 2010; Finocchio DV, et al., 1991; Gauthier GM and Robinson DA, 1975; Roucoux A, et al., 1983; Sherman KR and Keller EL, 1986). Purkinje cells in the flocculus reach mature levels in terms of activity level as well as dendritic and axonal morphology at three weeks after birth, and both the VOR and its adaptation already function virtually optimally at this early stage (Faulstich et al., 2004, Vis Res; Beekhof et al., 2021, Elife). In line with an early completion of the first process, here too we found that even at P18-20, less than one week after opening their eyes, both control and Purkinje cell specific PP2B knockout mice exhibited normal eye movement performance and learning. One week later, at age of P26-30, both basal reflexes, the VOR and the OKR, are affected by a deletion of PP2B. In contrast, in adult mice neither acute block, nor adult-onset prolonged deletion of PP2B affected the VOR gain, indicating the importance of development in setting the reflex level. Based on these findings, we postulate that the third to fourth postnatal weeks are the critical phases for the priming, or setting, of basal VOR, also referred to as the VOR baseline.

Why does the onset and fine-tuning of the compensatory eye movements take place at the third to fourth postnatal week? First, it should be noted that although the expression of Cre in Purkinje cells starts in the first postnatal week (Barski JJ, et al., 2000), our data indicate that PP2B is a stable protein that remains partially present after deletion of the gene for at least two more weeks. Given that eye opening occurs at postnatal 12-14 days in mice (Ko H et al., 2013), it is possible that our method precluded the possibility to observe effects of PP2B deletion at earlier stages (Goode CT et al., 2001; Weissman BM, et al., 1989). In humans, it has been suggested that early visual experience and the maturation of visual pathways are important to establish a setpoint for the VOR (Goode CT, et al., 2001). The morphological ontogenesis of the cerebellum, which is considered to be cerebellum-intrinsic and not input-dependent (Leto K, et al., 2016), could match that timeline and contribute to establishing the VOR setpoint (Weissman et al., 1989). The cerebellum undergoes its major growth in the third month and infant stage in humans, and the first 2 weeks after birth in mice, primarily due to expansion of granule cell progenitors (Dobbing J and Sands J, 1973; Rakic P and Sidman RL, 1970). In mice, although some parameters continue to change into adulthood, the circuit and its activity are largely established by the end of the third week (Beekhof GC, et al., 2021; Leto K, et al., 2016; van Welie I et al., 2011). For example, in terms of morphology, Purkinje cells are initially innervated by multiple climbing fibers with similar strengths in the first postnatal week, but from P9 to P17, climbing fibers successively undergo functional differentiation, dendritic translocation and elimination (Hashimoto K and Kano M, 2013; Watanabe M and Kano M, 2011). Moreover, *in vivo* electrophysiological studies in anesthetized and awake mice found that the firing rate of complex spikes increased sharply at 3 weeks of age, whereas the firing rate of simple spikes gradually increased until 4 weeks of age (Arancillo M et al., 2015; Beekhof GC, et al., 2021), which matches with the onset timing of the behavioral phenotypes in the present study.

Given that cerebellar development is more protracted than that of other brain regions (Bayer SA, et al., 1993), it appears to be more vulnerable for genetic or environmental disruptions, which could ultimately increase the risk for cerebellum-dependent behavior. That could be why many developmental disorders, such as autism, attention deficit hyperactivity disorder and developmental dyslexia, have been suggested to have cerebellar deficiencies (Leto K, et al., 2016; Manto MU and Jissendi P, 2012; Sathyanesan A et al., 2018; Wang R, et al., 2018; Wang SS, et al., 2014). Disruptions in Purkinje cell development can lead to deficits later on in life (Badura A et al., 2018; Sathyanesan A, et al., 2018), but the underlying principles are likely complex and currently far from clear. Therefore, it is key to elucidate how cerebellar development contributes to motor control in a systematic and comprehensive manner. Although numerous human studies have been performed to explore the developmental role of the cerebellum in eye movements (Alahyane N, et al., 2016; Charpiot A, et al., 2010; Finocchio DV, et al., 1991; Krishna O et al., 2018), rodent studies are overtly lacking. By directly examining the relationship between cerebellar development and compensatory eye movement at both the performance and adaptive functionalities, we were able to demonstrate that loss of cerebellar plasticity early in development has significant implications for basal motor behaviors at later stages, in that the third and fourth postnatal weeks seem particularly critical for the development of normal basal VOR function in mice.

Taken together, we here present a comprehensive, quantitative study into the effects of acute and (semi-)chronic disruption and recovery of plasticity at various life stages on the setpoint and adaptation of an innate reflex. The concept of a developmental, critical period to establish the baseline level of reflexes probably has important implications for sensorimotor functions in general. More work is needed to determine if and how our results can be converted to and possibly generalized across other modalities, including potential non-motor functions, and if so, whether it is possible to reset a deficit of developmental origin to normal levels through intervention during later stages.

## Acknowledgments

Financial support was provided by the National Natural Science Foundation of China (Grant # 82001202; BW), Shanghai Pujiang Talent Program (Grant # r 2020PJD006, BW), European Research Council (ERC-Stg #680235; MS).

## Appendix

